# Engineering and characterisation of AAVS1 FRT cell lines for inducible and dose-dependent gene expression

**DOI:** 10.64898/2026.10.29.762215

**Authors:** Maria Portillo Malumbres, Lachlan Ellingboe, Annalisa Roberti, Oxana Nashchekina, Fernando J. Sialana, Jonathon Pines, Jyoti S. Choudhary, Joerg Mansfeld, Norman E. Davey

## Abstract

Investigating biological function with high reproducibility requires the generation of stable cell lines that offer precise and inducible transgene expression. However, traditional methods often suffer from the limitations of random genomic integration, including unpredictable expression levels, leaky basal activity, and disruption of endogenous loci. To address these challenges, we engineered a versatile RPE1 cell line by integrating a TetOn3G-FRT cassette into the *AAVS1* safe-harbour locus using CRISPR/Cas9(D10A) nickase. This platform utilises the Flp-*FRT* recombination system to facilitate rapid, site-specific "plug- and-play" insertion of transgenes, combined with the Tet-On 3G system for tight, dose-dependent expression control. The resulting RPE1 *AAVS1* FRT8 line enables highly consistent transgene induction across independent clones, behaving as a monoclonal population. We highlight the importance of rigorous cell line characterisation through a multi-omics approach. Whole genome sequencing identified precise heterozygous integration at the target locus and *de novo* genomic duplications on other chromosomes. By integrating WGS with high-resolution proteomics and cell cycle analysis, we confirmed that these alterations did not significantly perturb cellular homeostasis or phenotypic fitness. This work establishes a robust and reliable platform for the efficient generation of inducible *AAVS1* FRT stable cell lines, facilitating reproducible research in modern molecular biology.

## Introduction

Human immortalised cells are central to modern biomedical research, providing a robust framework to investigate gene function, drug metabolism, cytotoxicity, and the production of vaccines and antibodies. A common experimental strategy to characterise biological processes involves the expression of exogenous, often epitope-tagged, proteins to enable biochemical, imaging, or functional analyses. The generation of such cell lines typically relies on transient or stable transfection. While transient transfection is rapid and technically straightforward, it is limited by poor reproducibility, heterogeneous expression across cell populations, and short-lived expression due to the lack of genomic integration. In contrast, cell lines generated via stable transfection offer sustained and more consistent expression, but their generation is more time- and labour-intensive, requiring selection of successfully transfected cells and extensive validation of stable expression.

Several strategies have been developed to facilitate the generation of stable transgene integration, including site-specific recombination-based methods such as Cre-LoxP (1) and Flp- *FRT* recombination (2) as well as transposon-based systems such as PiggyBac (3), and lentiviral transduction (4). Although widely used and effective, each approach has inherent limitations. PiggyBac transposition and lentiviral transduction result in random genetic integration, often leading to variable expression levels and potential disruption of endogenous genes , 6). Conversely, Cre-LoxP and Flp/*FRT* systems require prior introduction of recombination sites into the genome (7). Although these systems all enable subsequent targeted integration into a defined locus, thereby reducing expression variability, the initial integration of the recombination sites itself is often achieved through random integration. Consequently, the recombination sites may be inserted into genomic regions that are suboptimal for transgene expression. Clustered regularly interspaced short palindromic repeats (CRISPR)-based genome engineering offers a solution to this limitation facilitating the precise insertion of exogenous sequences at defined genomic loci (8). However, the generation of CRISPR-modified monoclonal cell lines remains resource-intensive, requiring substantial time, labour, and experimental resources for the design, screening, and validation of individual clones. Additionally, because genome engineering and clonal selection can introduce or enrich unintended genomic changes, validation of engineered cell lines should extend beyond confirmation of the intended insertion to include broader assessment of genomic integrity and cellular fitness. As a result, CRISPR-mediated targeted integration is often impractical when multiple cell lines expressing different constructs are required., 6). Conversely, Cre-LoxP and Flp/*FRT* systems require prior introduction of recombination sites into the genome (7). Although these systems all enable subsequent targeted integration into a defined locus, thereby reducing expression variability, the initial integration of the recombination sites itself is often achieved through random integration. Consequently, the recombination sites may be inserted into genomic regions that are suboptimal for transgene expression. Clustered regularly interspaced short palindromic repeats (CRISPR)-based genome engineering offers a solution to this limitation facilitating the precise insertion of exogenous sequences at defined genomic loci (8). However, the generation of CRISPR-modified monoclonal cell lines remains resource-intensive, requiring substantial time, labour, and experimental resources for the design, screening, and validation of individual clones. Additionally, because genome engineering and clonal selection can introduce or enrich unintended genomic changes, validation of engineered cell lines should extend beyond confirmation of the intended insertion to include broader assessment of genomic integrity and cellular fitness. As a result, CRISPR-mediated targeted integration is often impractical when multiple cell lines expressing different constructs are required.

A further critical consideration in generating engineered cell lines is the degree of precision required for controlling gene expression. There has been increasing interest in performing experiments at or near endogenous protein levels, reflecting the importance of maintaining physiologically relevant protein abundance for normal cellular function and homeostasis (9). Both excessive gene dosage and uncontrolled overexpression, particularly of dominant-negative or otherwise toxic protein products, can alter cellular processes and confound phenotypic interpretation, undermining physiological relevance of the models produced. While several constitutive promoters are routinely used for stable transgene expression, they offer no control over expression levels and are poorly suited for experiments requiring fine-tuning of gene dosage. Inducible and titratable expression systems provide a more appropriate framework for physiologically relevant experimental design, as they enable both temporal control and dose-dependent modulation of transgene expression (10). Among these, tetracycline (Tet)-dependent transcriptional systems are the most widely adopted (11). These systems typically consist of a Tet-responsive promoter and a transcriptional activator or repressor whose activity is regulated by tetracycline or its analogue doxycycline. However, despite their broad utility, earlier TetOn and TetOff implementations can suffer from basal leakiness and variable inducibility, often influenced by the genomic integration site of system components (12, 13). More recent systems, such as TaKaRa’s Tet-On 3G system, exhibit substantially reduced basal activity and enhanced sensitivity to doxycycline, allowing tight, inducible, and dosable control of transgene expression (14).

To combine the precision of CRISPR-targeted genome editing, the efficiency of Flp-*FRT* recombination, and the tight control of Tet-dependent expression, we sought to generate a human cell line containing an FRT recombination site and a Tet-On 3G transactivator (3G) integrated at the adeno-associated virus integration site 1 (*AAVS1*) safe-harbour locus using CRISPR/Cas9(D10A) nickase machinery (Figure 1A). Once a monoclonal *AAVS1* FRT cell line is characterised, subsequent insertion of exogenous constructs via Flp recombinase-mediated cassette exchange can be used to generate stable cell lines without further clonal selection. In this configuration, constructs of interest are cloned into a donor plasmid containing a TRE3GS inducible promoter and a compatible FRT site, allowing precise and reproducible integration into the *AAVS1* locus. The *AAVS1* locus, situated within the first intron of PPP1R12C on chromosome 19, was selected as the integration site because it is a well-established safe harbour that supports reliable transgene expression without perturbing endogenous gene function (15).

**Figure 1.**
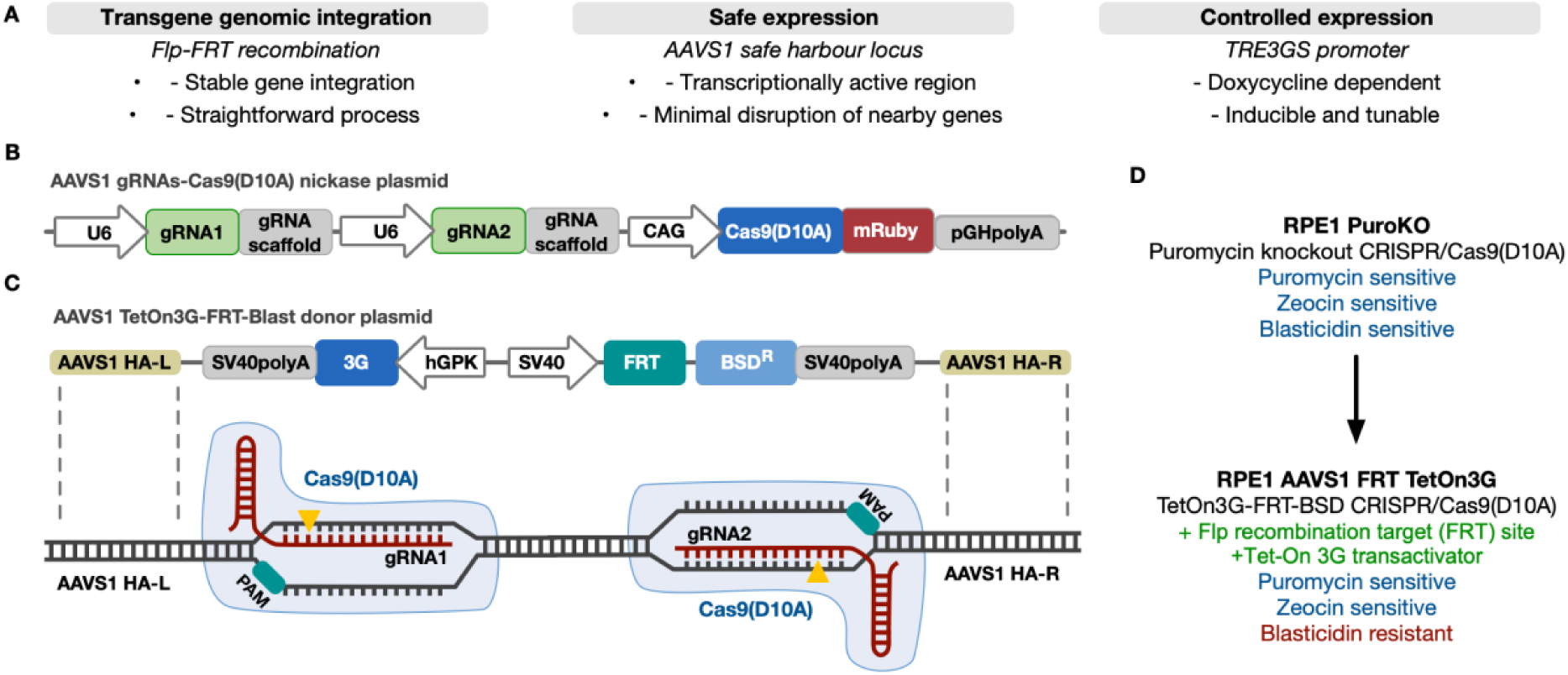
CRISPR/Cas9(D10A)n strategy for *AAVS1* FRT site knock-in in RPE1 cells. A) Features and advantages of an engineered *AAVS1* FRT TetOn3G cell line. B) Schematic of the Cas9(D10A) nickase plasmid. C) Donor plasmid carrying TetOn3G-FRT-BSD^R^ cassette flanked by *AAVS1* locus homology arms (HAs). D) Features of parental RPE1 cell line and engineered RPE1 *AAVS1* FRT TetOn3G cell line. Inserted expression modules are shown in green, antibiotic sensitivity in blue, and antibiotic resistance markers in red.

Together, the developed reagents and protocols for *AAVS1* FRT TetOn3G cell line engineering represent a versatile and efficient platform for the generation of stable, inducible cell lines with defined genomic integration. Here, we describe insertion at the *AAVS1* locus of RPE1 cells using CRISPR/Cas9(D10A) nickase. We detail the plasmid constructs, guide RNAs (gRNA), homology arms (HAs), and PCR-based genotyping strategies used for cell line generation and validation. Using this platform, we have generated an RPE1 *AAVS1* FRT TetOn3G cell line and validated the functionality of the Flp-*FRT* site-directed recombination system and the Tet-On 3G inducible gene expression system. We report successful integration of enhanced green fluorescent protein (eGFP) at the *AAVS1* locus and uniform, inducible, dose-dependent expression upon doxycycline addition. Finally, we provide a comprehensive genomic and proteomic characterisation of both the engineered RPE1 *AAVS1* FRT TetOn3G and the parental RPE1 cell line to validate the use of the engineered cell lines beyond simple confirmation of successful genetic modification.

## Results

### Design of AAVS1 gRNAs-Cas9(D10A)nickase plasmid and AAVS1 TetOn3G-FRT-BSD^R^ donor plasmid

To target the *AAVS1* locus, a plasmid containing a Cas9(D10A) nickase and gRNAs targeting two adjacent regions on opposite strands of the *AAVS1* locus was constructed (Figure 1B; Table S1). Both gRNA_HAL and gRNA_HAR were designed based on gRNA sequences previously validated by Bhagwan *et al.* (16) and Natsume *et al.* (17), respectively. Using a double-nicking strategy, both *AAVS1* gRNA-Cas9(D10A)n ribonucleoprotein complexes co-localised, bound, and nicked the *AAVS1* locus. Since the single-strand nicks were in close proximity, the result was a double-strand break. This double-strand break has long overhangs instead of blunt ends and is preferentially repaired by the homology-directed repair pathway in the presence of a donor construct (18). Ultimately, this reduces the probability of off-target insertion and leads to improved control of targeted insertions (19, 20).

As a donor construct, a plasmid containing an FRT site and Tet-On 3G transactivator flanked on each side with ∼800 bp of homology to the *AAVS1* locus was constructed (Figure 1C; Table S2). These HAs facilitate homologous recombination and ensure specific integration of the donor template into the *AAVS1* locus. Both the *AAVS1* gRNAs-Cas9(D10A)n plasmid and *AAVS1* TetOn3G-FRT donor plasmid were designed with selectable markers: gRNA-Cas9(D10A)n was co-expressed with mRuby to validate successful transfection and the donor plasmid contained a blasticidin resistance (BSD^R^) gene for clonal selection (Figure 1D).

### Engineering and genomic validation of RPE1 AAVS1 FRT TetOn3G cell line

Human hTERT-immortalised retinal pigment epithelial-1 (RPE1) cells, a genetically stable, non-transformed human cell line widely used to study physiological gene function was selected as the host cell line (21, 22). To achieve precise control over transgene expression, we sought to create a heterozygous RPE1 *AAVS1* FRT TetOn3G cell line in which only one *AAVS1* locus was modified by the integration of the TetOn3G-FRT-BSD^R^ cassette. The rationale for choosing heterozygous integration was to ensure that only one FRT recombination site was integrated into the genome, limiting the expression of any construct inserted via Flp-*FRT* recombination to arise specifically from a single site. To achieve this, RPE1 cells were co-transfected with the *AAVS1* gRNAs-Cas9(D10A)n plasmid and the *AAVS1* TetOn3G-FRT-BSD^R^ donor plasmid. Clones that had successfully integrated the TetOn3G-FRT-BSD^R^ cassette were isolated by screening for blasticidin resistance for 20 days followed by single cell sorting. Once the single cells had recovered and proliferated, genomic DNA from 101 clones was extracted for PCR genotyping using sets of primers spanning outer and inner regions of the integration site (Figure 2A).

**Figure 2.**
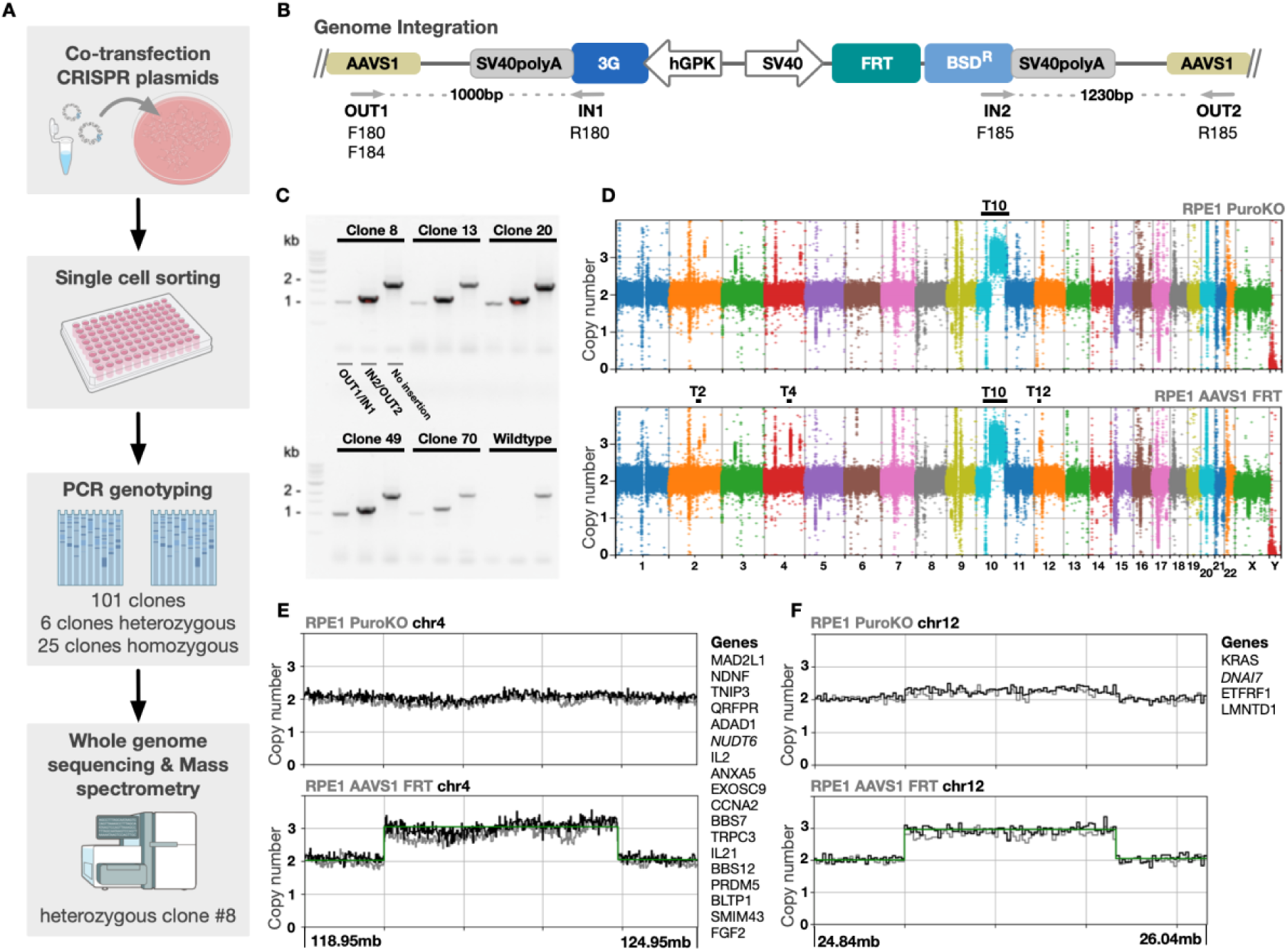
Engineering and genomic validation of FRT-site knock-in. A) Workflow to validate successful integration of TetOn3G-FRT-BSD^R^ cassette into the *AAVS1* locus. B) *AAVS1* locus after successful knock-in of TetOn3G-FRT-BSD^R^ cassette, with positions of PCR genotyping primers indicated in grey. C) Genomic DNA PCR screening of RPE1 *AAVS1* FRT TetOn3G clones #8, #13, #20, #49, and #70 with primers F180/R180, F185/R185, and F184/R185 (left to right). D) Manhattan plot of read depth across the whole genome for RPE1 PuroKO and RPE1 *AAVS1* FRT8. Generated by CNVPytor using a bin width of 10 Kbp. Duplication events T2, T4, T10, and T12 have been labelled. E, F) Read depth plots for the regions containing events T4 (E) and T12 (F). A list of genes covered by each duplication is to the right of each plot. Line colours; grey is uncorrected read depth, black is guanine-cytosine content corrected read depth, and green is estimated read depth.

To validate successful integration into the *AAVS1* locus, primer F185 was designed to bind to the TetOn3G-FRT-BSD^R^ cassette and R185 was designed to bind the *AAVS1* genomic locus downstream of the integration site to generate a 1230-bp PCR fragment. Primers F184/R185 were designed to bind upstream and downstream of the integration site to validate heterozygosity (Figure 2B, Table S3). In a wild-type locus, the PCR fragment would be 1980 bp in length, while a modified locus will yield a PCR fragment of 4533 bp. Using these two sets of primers, successful heterozygous integration of TetOn3G-FRT-BSD^R^ cassette in the *AAVS1* locus was confirmed for 6 of the 101 clones tested. An additional 26 colonies were confirmed for homozygous integration showing only the 1230-bp band (Figure S1). Of the six heterozygous clones, clones #8, #13, #20, #49, and #70 were further genotyped via PCR with an additional pair of primers (F180/R180) that bind upstream of the integration site and in the 3’ end of the cassette, resulting in a 1000-bp PCR fragment. These results confirmed the full integration of TetOn3G-FRT-BSD^R^ at the *AAVS1* locus and excluded the possibility of a truncated version (Figure 2C). PCR products from clones #8, #13, #20, #49, #70 were sequenced and no mutations were detected in the TetOn3G-FRT-BSD^R^ cassette.

We then extracted genomic DNA from RPE1 *AAVS1* FRT TetOn3G clone #8 (RPE1 *AAVS1* FRT8) and the parental cell line from which it was derived (RPE1 PuroKO) and performed whole genome sequencing (WGS). Sequencing was conducted using the Illumina NovaSeq600 platform, achieving a coverage depth of approximately 30x per genome. The data generated was analysed using SeqVerify, a recently developed computational pipeline designed to take raw WGS data, verify the presence of intended genomic edits, and detect any abnormalities or off-target effects (23). SeqVerify confirmed the insertion of the TetOn3G-FRT-BSD^R^ cassette into the *AAVS1* locus at chr19:58,209,44, cross-validating the results generated by PCR genotyping. No off-target insertions of TetOn3G-FRT were identified.

Additionally, the donor plasmid used for targeted insertion contained two A→T substitutions in each *AAVS1* HA to serve as markers of successful integration. These substitutions were detected in approximately half of the reads spanning the edited region. No reads containing these edits spanned the *AAVS1* locus, consistent with correct on-target integration (Figure S2). We identified several regions of trisomy: T2, T4, T10, and T12. T10 is a known feature of RPE1 cells, in which a segment of chromosome 10’s long arm is duplicated and translocated to the telomere of chromosome X (24). As expected, T10 was present both in RPE1 *AAVS1* FRT8 and the parental cell line (Figure 2D). In contrast, T2, T4 and T12 were unique to RPE1 *AAVS1* FRT8. T2 contained a single gene, *LRP1B*, whereas T4 contained 18 genes (Figure 2E). T12 spanned four genes and was trisomic in RPE1 *AAVS1* FRT8, while exhibiting a read depth of 2.29 in the parental cell line (Figure 2F), indicating a heterogeneous population, consisting of cells with and without the T12 duplication.

### RPE1 AAVS1 FRT TetOn3G preserves RPE1 cell cycle dynamics following CRISPR/Cas9(D10A) engineering

Following the identification of genomic duplications in the engineered RPE1 *AAVS1* FRT8 cell line, we investigated whether these alterations led to protein overexpression that might perturb cell cycle progression. This was of particular concern as the duplicated regions included *CCNA2* and *MAD2L1*, both of which are critical regulators of mitosis and the S/G2 transition (25, 26). To determine if these duplications translated to protein abundance changes, we performed whole proteome analysis by quantitative mass spectrometry. Of the 23 duplicated genes, eight corresponding proteins were detected, including MAD2L1, CCNA2, and KRAS. Both MAD2L1 and CCNA2 had an increase in abundance proportional to their increase in copy number, whereas KRAS had an unexpected decrease in abundance (Figure 3A, Figure S3).

**Figure 3.**
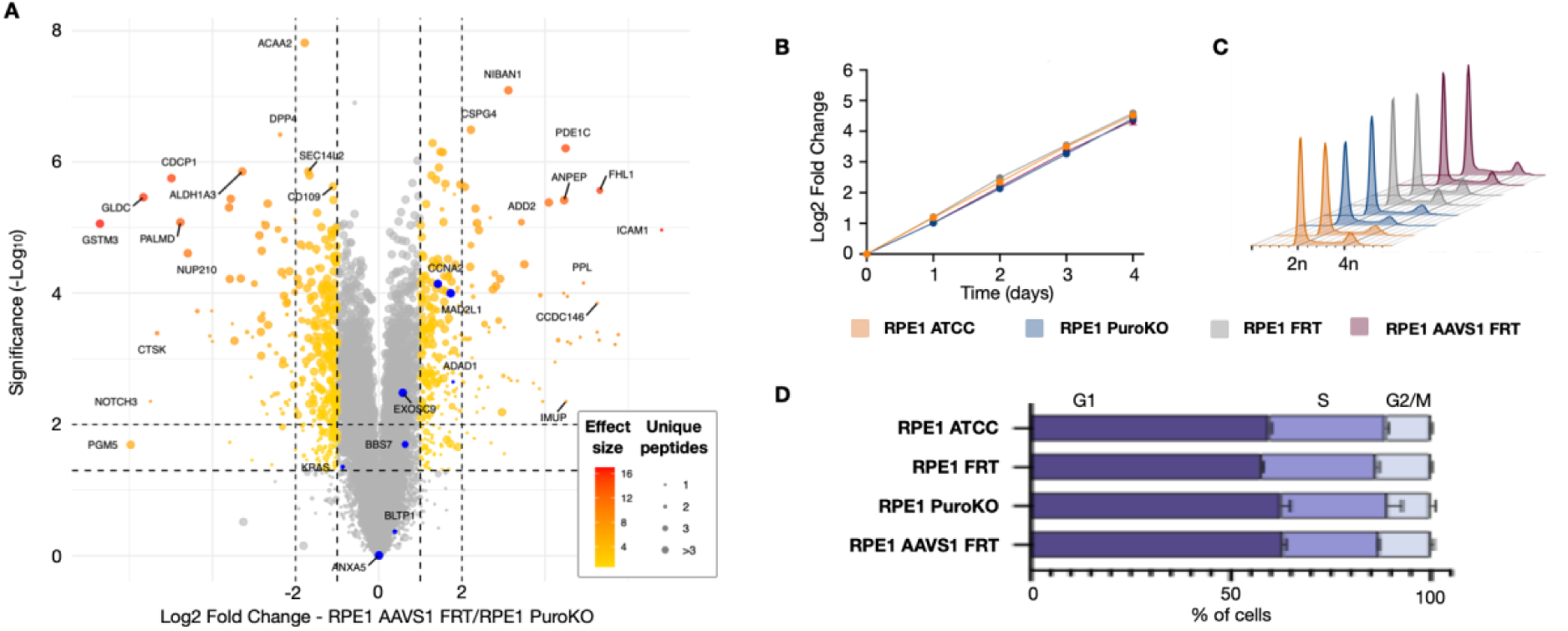
Characterisation of RPE1 *AAVS1* FRT TetOn3G proteome and cell cycle dynamics. A) All proteins identified by quantitative mass spectrometry (n=3 biological replicates). The top 20 hits by effect size are labelled and the eight proteins originating from a duplicated gene are both labelled and coloured blue. Horizontal dashed lines are drawn at p-values of 0.05 and 0.01, vertical dashed lines are drawn at fold change values of ±0.5 and ±1. B) Comparison of RPE1 ATCC, RPE1 puromycin knockout (PuroKO), RPE1 FRT, and RPE1 *AAVS1* FRT8 cell lines’ proliferation rate via cell viability assay. Luminescent signal was measured every 24 h to determine ATP levels by addition of CellTiter-Glo reagent (Promega). Luminescence intensity at each timepoint was converted to doublings relative to time zero for plotting. Points are means of triplicate wells. C) Peaks correspond to DNA content, with the z-axis separating samples for clarity. Each profile corresponds to an independent sample acquired under identical flow cytometry settings. D) Cell cycle distribution of RPE1 ATCC, PuroKO, FRT, and *AAVS1* FRT cell lines, shown as the average percentage of cells in G1, S, and G2 phases across two biological replicates. Cell-cycle phase distributions were normalised to 100% for each cell line. Statistical analysis was performed using two-way repeated-measures ANOVA followed by Dunnett’s multiple-comparisons test, comparing each cell line with RPE1 ATCC within each cell-cycle phase. No statistically significant differences were detected.

We further examined the proliferation rate of RPE1 *AAVS1* FRT8 and compared it to parental RPE1 cell lines. We included the original RPE1 cell line from American Type Culture Collection (ATCC), the parental RPE1 puromycin knockout cell line (27) used to generate RPE1 *AAVS1* FRT8 cell line, and an additional RPE1 FRT cell line in which an FRT site was randomly integrated (28) in this comparative analysis. To determine the number of viable cells in culture, a CellTiterGlo luminescence assay was used to quantify ATP, an indicator of metabolically active cells (29). To assess proliferation without cell passaging, 400 cells were seeded per well. Time zero was measured 6 h after seeding and additional time points were measured every 24 h for 4 d. All four RPE1 cell lines showed a comparable proliferation profile, doubling approximately every 24 h (Figure 3B). To assess whether the genetic manipulation induced any abnormal cell cycle behaviour, we performed cell cycle profiling by flow cytometry using DAPI staining as a marker of DNA content across the RPE1 *AAVS1* FRT8 and the other RPE1 cell lines included in the proliferation analysis (Figure 3C). The DAPI fluorescence intensity profiles were then used to quantify the cell cycle distribution. The percentage of cells in the G1, S, and G2/M phases for each condition was calculated and averaged over two replicates (Figure 3D). Comparable cell cycle distributions were observed in all cell lines.

### RPE1 AAVS1 FRT8 cell line enables dose-dependent control of transgene expression after Flp/FRT recombination

Having confirmed that the RPE1 *AAVS1* FRT8 TetOn3G cell line maintains normal cellular physiology, we next validated the functionality of the Flp-*FRT* site-directed recombination system and the TRE3GS inducible promoter (Figure 4A and 4B). We constructed an FRT expression plasmid encoding the TRE3GS promoter, eGFP, and neomycin resistance gene (TRE3GS-eGFP-FRT-Neo^R^) (Figure 4C). This expression plasmid was co-transfected with a flippase helper plasmid into the RPE1 *AAVS1* FRT8 cell line. Successful site-specific Flp/FRT recombination into the *AAVS1* locus places the Neo^R^ gene under the control of the constitutive SV40 promoter, allowing for the positive selection of clones, while eGFP expression remains under the control of the doxycycline-inducible TRE3GS promoter (Figure 4D).

**Figure 4.**
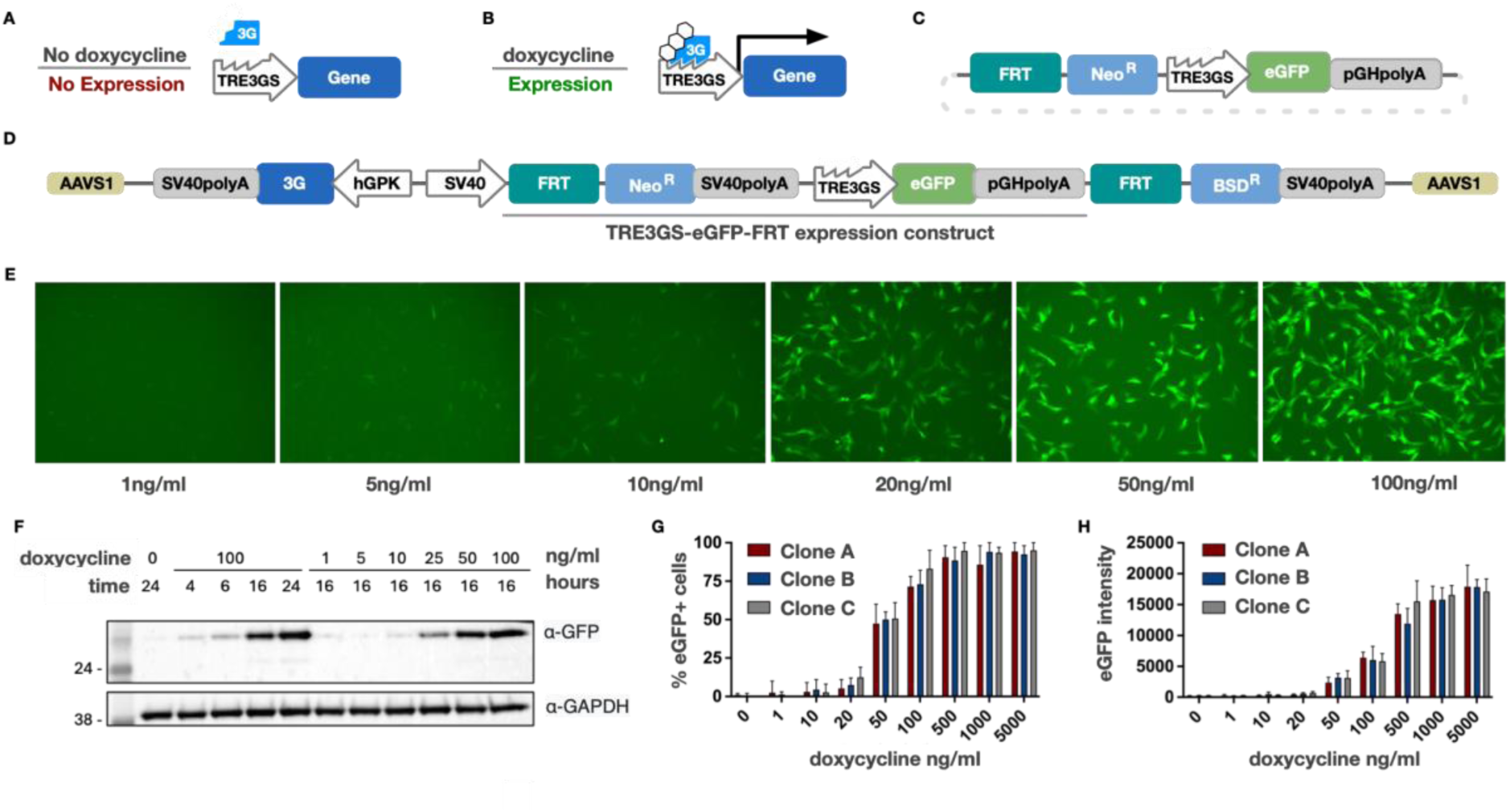
Validation of Flp/FRT site-directed recombination and TetOn3G system inducible gene expression. A, B) Schematic representation of the Tet-On 3G induction mechanism. In the absence of doxycycline (A), the Tet-On 3G transactivator cannot bind the TRE3GS promoter. Upon addition of doxycycline (B), the transactivator undergoes a conformational change, binds to TRE3GS, and initiates transgene expression. C) Schematic of the FRT-expression plasmid containing the TRE3GS-eGFP-FRT-Neo^R^ cassette. D) Genomic architecture of the RPE1 *AAVS1* locus following successful Flp/FRT-mediated integration. The Neo^R^ gene is placed under the control of the endogenous SV40 promoter, while eGFP is regulated by the inducible TRE3GS promoter. E) Representative fluorescence microscopy images of RPE1 cells showing eGFP expression 16 h post-induction with increasing concentrations of doxycycline (1–100 ng/ml). F) Western blot analysis of eGFP protein levels. Left: Time-course of eGFP induction following treatment with 100 ng/ml doxycycline. Right: Dose-response analysis of eGFP levels across a doxycycline gradient at 16 h post-induction. GAPDH serves as the loading control. Data are representative of three independent biological replicates (n=3). G) Quantification of the percentage of eGFP-positive cells via live-cell imaging for three independent clones (A, B, and C) across a broad doxycycline concentration range (0–5000 ng/ml). Bars represent the mean of two independent experiments, with duplicate measurements per experiment (n=2). H) Mean eGFP fluorescence intensity for Clones A, B, and C across the indicated doxycycline gradient. Bars represent the mean of two independent experiments, with duplicate measurements per experiment (n=2).

Following 10 days of selection with G418 (analog of neomycin), cells were induced with 1 µg/ml doxycycline and eGFP-positive cells were isolated via fluorescence-activated cell sorting (FACS; Figure S4). The resulting population was expanded and treated with increasing concentrations of doxycycline, revealing a robust dose-dependent increase in eGFP expression (Figure 4E). These findings were corroborated by western blot analysis, which detected eGFP as early as 4 h post-induction, with protein levels continuing to accumulate over a 24-h period (Figure 4F).

To evaluate the robustness and clonal consistency of the system, we isolated three independent clones (FRT8 A, B, and C) via single-cell sorting. We then quantified both the percentage of eGFP-positive cells and the mean fluorescence intensity using live-cell imaging. At 50 ng/ml doxycycline, approximately 50% of the population showed detectable eGFP expression (Figure 4G). However, at concentrations of 100 ng/ml and above, the response became more uniform, with 80%–90% of cells expressing eGFP and exhibiting a dose-dependent increase in fluorescence intensity (Figure 4G and 4H). The consistency among independent clones confirms that the RPE1 *AAVS1* FRT8 platform yields monoclonal-like reliability for controlled doxycycline-dependent transgene expression.

## Discussion

Studying biological function in a reproducible manner usually requires the generation of stable cell lines that express exogenous transgenes with high fidelity. With this work, we provide the molecular reagents and the framework to engineer and validate a TetOn3G-FRT cassette into the *AAVS1* safe harbour locus using a CRISPR/Cas9(D10A) nickase, avoiding the common pitfalls associated with random integration, such as unpredictable expression levels, leaky basal activity, and disruption of endogenous genes.

The AAVS1 locus, within intron 1 of *PPP1R12C*, is a well-established safe harbour (15). Its open chromatin supports stable, robust transgene expression with minimal silencing, insertions there are well tolerated by cells, and it is efficiently targeted by CRISPR/Cas9. Integrating at a defined site also removes the position-effect variability of random integration, improving reproducibility between lines. AAVS1 is therefore widely used for targeted integration. For example, a recent study introduced inducible Cre-loxP systems at this locus in RPE1 cells (30).

In this study, we introduce a Flp–*FRT* recombination system for genomic integration to facilitate stable, site-specific insertion through a streamlined workflow. This process is highly compatible with "plug-and-play" applications, allowing researchers to generate multiple cell lines expressing different transgenes in parallel from a single master line. We chose to use the Tet-On 3G (TaKaRa) system due to its significant improvements over previous generations. This mechanism ensures tight, inducible, and tuneable control over transgene expression levels.

While the AAVS1 gRNA-CRISPR/Cas9(D10A) and TetOn3G-FRT-BSD^R^ donor plasmids were validated specifically in RPE1 cells, we anticipate their broad utility for targeting this safe-harbour locus across diverse human cell lines. Furthermore, both the donor and expression plasmids are highly modular; the latter can be easily adapted for the Flp-mediated integration of any transgene of interest. By sharing these reagents and protocols, we seek to provide the broader research community with a streamlined path toward efficient, site-specific genome editing at the *AAVS1* locus. To facilitate the adoption of this platform, the following plasmids have been deposited with Addgene: *AAVS1* gRNAs-Cas9(D10A)n plasmid (#252799); *AAVS1* TetOn3G-FRT-BSD^R^ donor plasmid (#252870); TRE3GS-eGFP-FRT expression plasmid (#252871).

Additionally, the engineered RPE1 *AAVS1* FRT8 TetOn3G master cell line is being prepared for deposition at CancerTools.org, with the aim of providing a ready-to-use host for inducible transgene expression.

After designing and engineering of the RPE1 *AAVS1* FRT TetOn3G cell line, we validated the system’s functionality by demonstrating the inducible and dose-dependent expression of eGFP upon treatment with doxycycline. While the Flp-*FRT* recombination procedure is straightforward, we observed that transfection efficiency in RPE1 lines remained a bottleneck, typically ranging from 5%–30%, even under optimised electroporation conditions. Expanding this TRE3GS-FRT system to other human cell lines with higher transfection efficiencies will likely enhance the success rate of generating stable, inducible cell lines.

A critical takeaway of this work is the necessity of performing validations beyond confirming the intended genomic modification. As CRISPR-mediated engineering can result in unintended genomic alterations, we advocate for the adoption of WGS and mass spectrometry as standard practice in cell line development. With the recent reduction in sequencing costs and the availability of streamlined bioinformatic pipelines, these analyses represent a relatively small additional cost compared with the substantial investment required to generate a cell line, particularly when considered against the risk of proceeding with an inadequately characterised model. In this study, WGS enabled us to pinpoint the precise integration coordinates at chr19:58,209,44 and confirm the heterozygosity of the insertion. Furthermore, WGS identified *de novo* gene duplications on other chromosomes. By integrating WGS with whole-proteome analysis and viability assays, we confirmed that these duplications did not observably perturb the cellular phenotype, ensuring the line remained "fit for purpose" as a research platform. However, we note that researchers intending to study specific pathways directly related to these duplicated genes should proceed with caution.

In summary, by combining site-specific integration with a tightly controlled inducible system, *AAVS1* FRT TetOn3G cell lines can serve as a versatile and reliable platform for the efficient generation of stable and inducible cell lines.

### Data Availability

The mass spectrometry proteomics data are available via ProteomeXchange in the PRIDE repository under accession PXD085689 (https://www.ebi.ac.uk/pride/archive/projects/

PXD085689). Whole-genome sequencing data are available from the corresponding author upon request.

## Acknowledgements

We would like to thank Danny D’Amore for detailed editing of the manuscript. We thank the ICR facilities for their support with cell sorting and imaging. We thank the Anastassiadis Lab (Dresden University of Technology) for the gift of the plasmids for the AAVS1 homology arms.

N.E.D. and M.P.M are funded by a Cancer Research UK Senior Cancer Research Fellowship (C68484/A28159). N.E.D. is funded by the American Lebanese Syrian Associated Charities.

## Methods

### AAVS1 gRNAs oligo design

gRNA_HAL and gRNA_HAR oligos were designed based on gRNA sequences previously validated by Bhagwan *et al.* (16) and Natsume *et al.* (17). For both oligo pairs, protospacer adjacent motif (PAM) sites were removed and BbsI or BsaI overhangs were added. The final oligos (Table S1) were ordered from Integrated DNA Technologies at 25 nM with Standard Purification. gRNA_HAL and gRNA_HAR oligos were annealed by mixing 20 µl of 100 µM oligo forward and reverse primers and heating at 95°C for 5 min before allowing the mixture to gradually cool to ambient temperature.

### AAVS1 TetOn3G-FRT donor plasmid cloning

The *AAVS1* TetOn3G-FRT donor plasmid was generated by insertion of a pSV40-FRT-SV40pA fragment synthesised by Genewiz into a pDrive backbone using T4 Quick Ligase (New England Biolabs; NEB). The blasticidin resistance gene (BSD^R^) was PCR amplified using the Expand High Fidelity PCR System (Roche) per the manufacturer’s instructions and inserted into pDrive-pSV40-FRT-SV40pA plasmid. *AAVS1* left and right HAs (Table S2) were PCR amplified from a plasmid kindly provided by the Anastassiadis Lab (Dresden University of Technology) with primers designed to mutate the PAM sequence before being inserted sequentially into the pDrive-pSV40-FRT-BSD^R^-SV40pA plasmid. hPGK-T3G transactivator-SV40pA was PCR amplified using Phusion HF DNA Polymerase (NEB) from the TaKaRa Tet-One™ Inducible Expression System and ligated into the pDrive-HAL-pSV40-FRT-BSD^R^-SV40pA-HAR plasmid. Ligation of the PCR products was performed using the NEBuilder HiFi DNA Assembly Master Mix (NEB). Successful assembly of each intermediate plasmid and final plasmid was verified by Sanger sequencing.

### Cell culture

RPE1 ATCC (CRL-4000), RPE1 puromycin KO (kindly provided by Steve Jackson, Cancer Research UK Gurdon Institute), RPE1 FRT (kindly provided by Jonathon Pines, The Institute of Cancer Research, London), and RPE1 *AAVS1* TetOn3G FRT cells were cultured in Dulbecco’s Modified Eagle Medium F12 (DMEM:F12, Sigma Aldrich) supplemented with 10% foetal bovine serum (FBS, Thermo Fisher Scientific), 7.5% sodium bicarbonate (Thermo Fisher Scientific), GlutaMAX (Thermo Fisher Scientific), and penicillin-streptomycin (Thermo Fisher Scientific). Cells were maintained in a humidified incubator at 37°C and 5% CO2 concentration.

### Transfection

RPE1 puromycin KO cells co-transfection with the *AAVS1* gRNAs-Cas9(D10A)n plasmid and *AAVS1* TetOn3G-FRT-BSD^R^ donor plasmid was conducted using Lipofectamine 3000 Reagent (Invitrogen). Cells were seeded in 15-cm dishes at 1e6 cells per dish. The next day, the cells were transfected with 5 ug DNA of each plasmid following the manufacturer’s instructions. Seventy-two hours after transfection, media was replaced and supplemented with 10 μg/ml blasticidin (Life Technologies) for positive selection of clones for up to 20 d. Media was replaced every 4 days to add fresh blasticidin. Co-transfection of RPE1 *AAVS1* FRT8 cells with flippase helper plasmid pOG44 and TRE3GS-eGFP-FRT expression plasmid was conducted using the Neon^TM^ Transfection System (Thermo Fisher Scientific). RPE1 *AAVS1* FRT8 cells were trypsinised and washed with PBS before 1e6 cells were transfected using a flippase:expression plasmid ratio of 5:1 for a total of 5 μg DNA. Cells were electroporated at 1400 V, 20 ms, for 2 pulses. Forty-eight hours after electroporation, cells were placed under positive selection with 400 μg/ml Geneticin™ Selective Antibiotic (G418 Sulfate; Thermo Fisher Scientific), which was maintained for up to 15 d. Cells were regularly passaged to avoid over-confluency, and selective medium was replaced every 4 days with fresh medium containing 400 μg/ml G418 Sulfate.

### Screening of RPE1 AAVS1 FRT positive clones

Following blasticidin selection, resistant RPE1 *AAVS1* TetOn3G FRT cells were single-cell sorted using a BD FACSymphony™ S6 cell sorter (BD Biosciences) to ensure monoclonality. Cells were detached with trypsin, washed with PBS, resuspended in Leibovitz’s L-15 Medium (L15; Thermo Fisher Scientific) supplemented with 10% FBS for a final concentration of 2e6 cells/ml, and filtered using a 30-μm cell strainer (Sysmex). Single cells were sorted into 96-well plates with media supplemented with 20% FBS to improve cell recovery. After 10 d, genomic DNA from 101 single clones was extracted for genotyping using Taq DNA Polymerase (QIAGEN) and the primers given in Table S3. Integration of the TetOn3G-FRT-BSD^R^ cassette into *AAVS1* locus was confirmed with primers F180/R180 and F185/R185. Unperturbed *AAVS1* loci were detected with primers F184/R185. For genomic DNA extraction, clones were trypsinised, resuspended in DirectPCR Lysis Reagent (PEQlab) with proteinase K (Sigma-Aldrich), and heated for 1 h at 55°C before being heated to 95°C for 50 min. Then, 200 ng genomic DNA was extraged and used for PCR amplification. PCR products were analysed by agarose gel to check efficient amplification. PCR cycle parameters were: 94°C for 5 min for initial denaturation; 4 cycles of denaturation at 94°C for 15 s, 63°C for 30 s, and 72°C for 135 s; 4 cycles of denaturation at 94°C for 15 s, 60°C for 30 s, and 72°C for 135 s; 40 cycles of denaturation at 94°C for 15 s, 57°C for 30 s, and 72°C for 135 s, with a final elongation step of 72°C for 10 min. PCR products from positive clones (#8, #13, #20, #49, and #70) were purified from the gel and further validated by Sanger sequencing.

### FACS

Cells were treated with 1 μg/ml doxycycline hyclate (Sigma-Aldrich) to induce expression from the TRE3GS promoter. RPE1 *AAVS1* FRT8 eGFP cells were detached with trypsin, washed once with PBS, and resuspended in L15 with 10% FBS for a final concentration of 2e6 cells/ml. Resuspended cells were filtered using a 30-μm cell strainer (Sysmex). Flow cytometry samples were sorted using a BD FACSymphony™ S6 cell sorter (BD Biosciences). First, single cells were gated based on forward scatter and side scatter to exclude doublets and debris. From the single-cell population, gates were applied to identify eGFP-positive cells. A total of 40,000 cells were sorted into 6-well plates containing pre-warmed DMEM:F12 media. Flow cytometry data were acquired using BD FACSDiva™ software version 9.0 (Becton Dickinson) and analysed with FlowJo version 10.9 (BD Biosciences).

### Cell Cycle Profiling

Cells were cultured as described above and detached from culture dishes by trypsinization. Cells were washed with PBS and fixed with 70% ethanol for 4 h at −20°C. Following fixation, cells were stained for 20 min with 1 µg/ml propidium iodide supplemented with 10 µg/ml RNase (Sigma). Stained cells were analysed using an LSR II flow cytometer (BD Biosciences). Cell cycle profiles were analysed using FlowJo version 10.9 (BD Biosciences). Statistical analyses were performed using GraphPad Prism. Cell cycle distribution was analysed by two-way repeated-measures ANOVA followed by Dunnett’s multiple-comparisons test, comparing each cell line with RPE1 ATCC within each cell-cycle phase. Data were obtained from two independent biological experiments.

### Cell viability assay

Cells were plated in flat bottom, chimney style, black μclear 96-well plates (Greiner Bio-One) at 400 cells/well. Time zero was measured 6 h after seeding. Every 24 h, cells were lysed using CellTiter-Glo 2.0 Luminescent Cell Viability Assay (Promega), and luminescence was measured using the SpectraMax iD3 (Molecular Devices). Fold change was calculated by normalising the luminescence values at each timepoint to the median value of time zero and expressed as log2 fold change. Each condition was measured in triplicates.

### Live cell imaging

To assess eGFP expression, fluorescence imaging was performed using an EVOS™ M5000 Imaging System (Thermo Fisher Scientific). A total of 600,000 RPE1 cells were seeded in a 10-cm Petri dish 24 h prior to treatment with increasing concentrations of doxycycline over 16 h. Images were captured at 10x magnification using the EVOS LED Cube, GFP 2.0 (Invitrogen). To quantify eGFP expression, automated microscopy was performed on an ImageXpress Micro XLS wide-field screening microscope (Molecular Devices) equipped with 10x, 0.5 NA and 20x, 0.7 NA Plan Apo air objectives (Nikon) and laser-based autofocus. Cells were plated in 96-well clear-bottom plates (μclear, Greiner Bio-One) at 3,000 cells/well 24 h before imagining. Cells were treated with different concentrations of doxycycline for 16 h. One hour before imaging, a 50-nM final concentration of Sir-DNA stain (Spirochrome) was added to each well. Cells were imaged in L15 media supplemented with 10% FBS and 1% penicillin/streptomycin. Image analysis was performed using automated pipelines developed in the Custom Module Editor of MetaXpress (Molecular Devices). Images were flat-field and background corrected using a top-hat filter. Nuclei were segmented based on Sir-DNA staining to generate nuclear masks, which were used to extract fluorescence intensities for eGFP. Positive cells were identified by applying fixed thresholds to flat-field–corrected GFP channels based on visual inspection. All conditions and cell lines were processed in parallel using identical thresholds.

### SDS-PAGE and western blotting

Cells pellets were lysed in extraction buffer (1% (v/v)) Triton X-100, 10 mM Tris-HCl, 5 mM EDTA, 50 mM NaCl, 0.1% NP-40, and 1mM PMSF, supplemented with Complete Mini Protease Inhibitor Cocktail (Roche) by passage through a 26-gauge needle six times. The lysate was incubated on ice for 5 min and then clarified by centrifugation at 14,000 rpm for 10 min at 4°C. Protein concentration was quantified with a Bradford assay. Proteins were resolved by SDS-PAGE and transferred onto a nitrocellulose membrane for immunoblot analysis. After 1 h blocking with 5% milk in TBST (Thermo Fisher Scientific) at room temperature for 1 h, membranes were incubated overnight at 4°C with GFP (Roche) and GAPDH (Santa Cruz) primary antibodies. After washing with TBST three times, membranes were incubated for 1 h at room temperature with anti-mouse IgG HRP (Cell Signalling) secondary antibody. Membranes were again washed three times with TBST and blots were detected with Pierce ECL Western Blotting Detection Reagent (Thermo Fisher Scientific) on an ChemiDocTM MP Imaging System (Bio-Rad).

### Mass Spectrometry

#### Cell Culture and Sample Preparation for Proteomics

For quantitative proteomic analysis, RPE1 ATCC, RPE1 PuroKO, RPE1 FRT, and RPE1 *AAVS1* FRT cell lines were analysed in biological triplicate. Cells (2×10^^6^) were harvested, washed twice with ice-cold PBS, snap-frozen, and stored at −80 °C. Samples were processed using the SimPLIT workflow as previously described. Briefly, cell pellets were lysed in 0.1 M TEAB containing 1% sodium deoxycholate (SDC), 10% isopropanol, 50 mM NaCl, 5 mM TCEP, 10 mM iodoacetamide, universal nuclease, and protease/phosphatase inhibitors. Following sonication and incubation at room temperature, protein concentrations were determined using the Bradford assay (Bio-Rad). Aliquots containing 25 μg protein were digested with trypsin (Pierce) and labelled with TMTpro reagents (Thermo Fisher Scientific) according to the manufacturer’s instructions.

Labelled peptides were combined in equal amounts, acidified with formic acid to precipitate SDC, and clarified by centrifugation. The resulting peptide mixture was dried by vacuum centrifugation prior to high-pH fractionation and LC-MS/MS analysis.

#### High-pH Reversed-Phase Fractionation

Peptides were fractionated by offline high-pH reversed-phase chromatography on an XBridge C18 column (2.1 × 150 mm, 3.5 μm; Waters) using an Ultimate 3000 HPLC system (Thermo Fisher Scientific). A total of 96 fractions were collected and concatenated into 24 pooled fractions for LC-MS/MS analysis.

#### LC-MS/MS Analysis

TMTpro-labelled peptides were analysed using an Orbitrap Ascend Tribrid mass spectrometer coupled to a Vanquish Neo UHPLC system (Thermo Fisher Scientific). Peptides were separated at a flow rate of 300 nL min^-1 using a 120 min gradient from 5% to 35% solvent B over 100 min, followed by washing at 95% solvent B and re-equilibration at 5% solvent B. MS^1 spectra were acquired in the Orbitrap (m/z 400-1600) at a resolution of 120,000 with an AGC target of 4 × 10^5. Precursors (charge states 2-6) were isolated using a 0.7 m/z quadrupole window and fragmented by HCD (NCE 32%), with MS^2 spectra acquired in the ion trap operating in turbo scan mode. Real-time database searching was performed against a human protein database with carbamidomethylation (C) and TMTpro labeling (peptide N-termini and K) specified as static modifications and methionine oxidation and N/Q deamidation as variable modifications. Quantitative SPS-MS^3 spectra were acquired using up to 10 SPS ions, HCD fragmentation (NCE 55%), and Orbitrap detection at 45,000 resolution. TMTpro reporter ions were quantified from SPS-MS^3 spectra.

#### Database Search and Quantification

Raw data were processed in Proteome Discoverer v2.4.0.305 (Thermo Fisher Scientific) using the SEQUEST HT search engine and searched against the UniProt Swiss-Prot human proteome supplemented with common contaminants. Search parameters included a precursor mass tolerance of 20 ppm, fragment mass tolerance of 0.5 Da, trypsin specificity with up to two missed cleavages, static modifications of carbamidomethyl (C) and TMTpro (N-termini/K), and variable modifications of methionine oxidation and asparagine/glutamine deamidation. Peptide-spectrum matches were filtered to 1% FDR using Percolator. TMTpro quantification was performed from SPS-MS^3 spectra using a 15 ppm reporter ion integration tolerance with isotopic impurity correction applied. Only unique peptides were used for protein quantification, requiring an average reporter ion signal-to-noise ratio >3 and ≥50% SPS mass match. Protein abundances were median-normalized across samples prior to downstream analysis

### WGS

#### Sample Preparation

Genomic DNA from RPE1 ATCC, RPE1 puromycin KO, RPE1 FRT, and RPE1 *AAVS1* FRT cell lines was extracted using the Wizard® SV Genomic DNA Purification System (Promega) following manufacturer’s instructions and quantified using DeNovix DS-11 spectrophotometer (DeNovix) to meet GENEWIZ’s requirements of >1.5 µg at >20 ng/µL. Samples were sent to GENEWIZ for library preparation and sequencing on Illumina NovaSeq6000 yielding a sequencing depth of approximately 30x per genome.

#### Insertion and Copy Number Variation Analysis

Paired-end raw FASTQ files were first processed using the target-validation software SeqVerify

(23). The insertion site in the reference genome was defined by analysing chimeric and non-chimeric read mapping. First, CNVPytor (31) was used within SeqVerify to identify copy number varitation (CNV) events. Then, reads were filtered with Sambamba (32) with the following options: <4 mismatches; alignment score >60; and map quality >40. CNVPytor was rerun at 10,000-bp resolution on the filtered reads. Novel CNV events were then determined by comparing each cell line to its parental cell line. To ensure the novel events were of high quality, local read depth plots were generated for each novel CNV event using RPE1 *AAVS1* FRT8 and its parental cell line. These plots were manually checked for a consistent read depth throughout the deleted/duplicated region. To generate Manhattan plots, CNVPytor’s interactive plotting environment was used. All plots were made using the Sambamba-filtered bam files. For the whole genome plot, a resolution of 10,000 bp was used with CNVPytor’s Manhattan styling. For local read depth plots, a resolution of 10,000 bp was used with CNVPytor’s default styling. The graph output was restricted to the deleted/duplicated region ±30% of the region’s size.

## Supplementary figures

**Figure S1.**
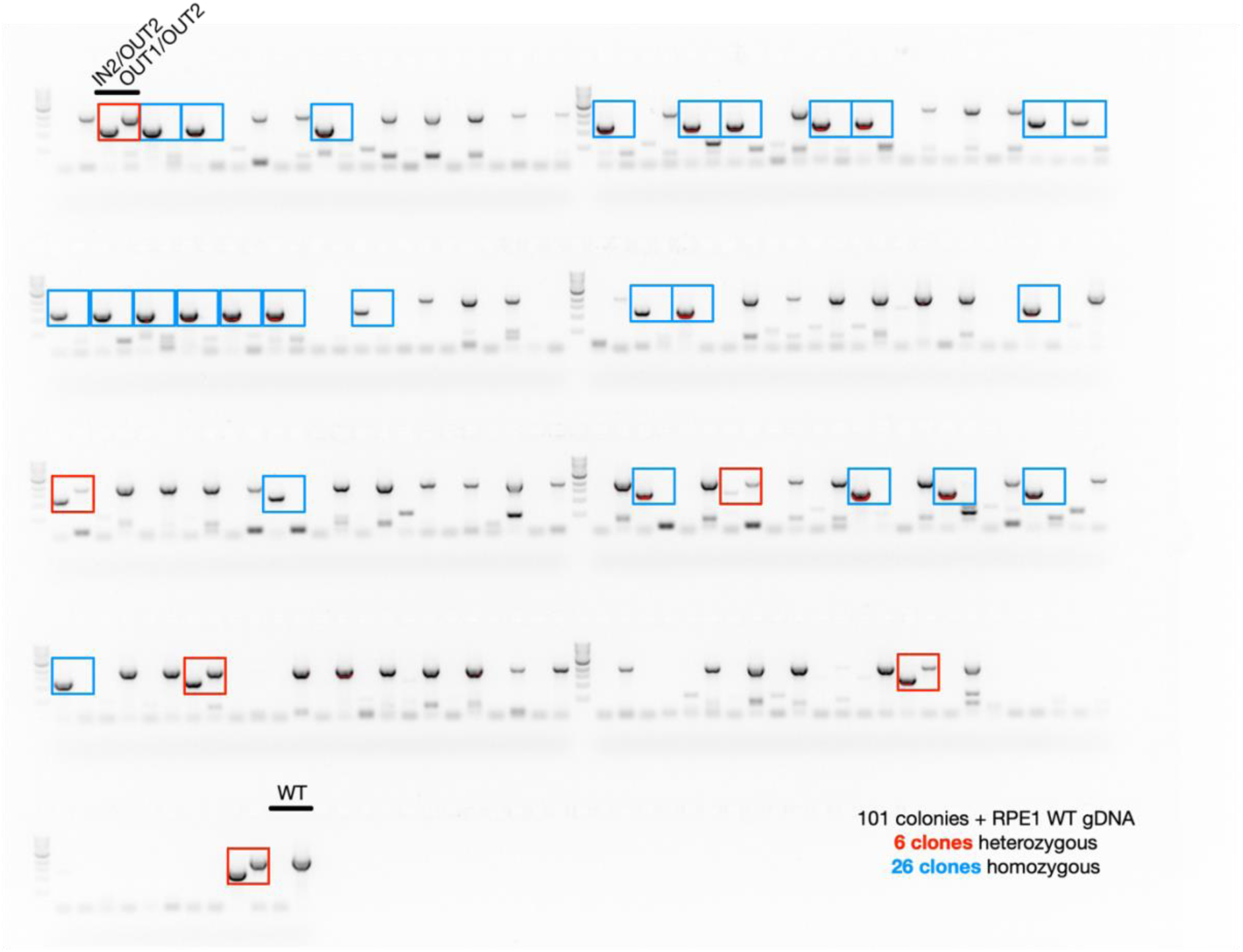
PCR genotyping of the RPE1 *AAVS1* locus. Related to Figure 2. Genomic DNA (gDNA) PCR screening of 101 RPE1 blasticidin resistant colonies after CRISPR/Cas(D10A)n engineering. Genomic DNA from wild-type (WT) RPE1 was included as negative control. IN2/OUT2 and OUT1/OUT2 primers correspond to F185/R185 and F184/R185, respectively. Colonies showing the expected band sizes for heterozygous TetOn3G-FRT-BSD^R^ integration are highlighted in red, while colonies showing the expected band sizes for homozygous TetOn3G-FRT-BSDR integration are highlighted in blue.

**Figure S2.**
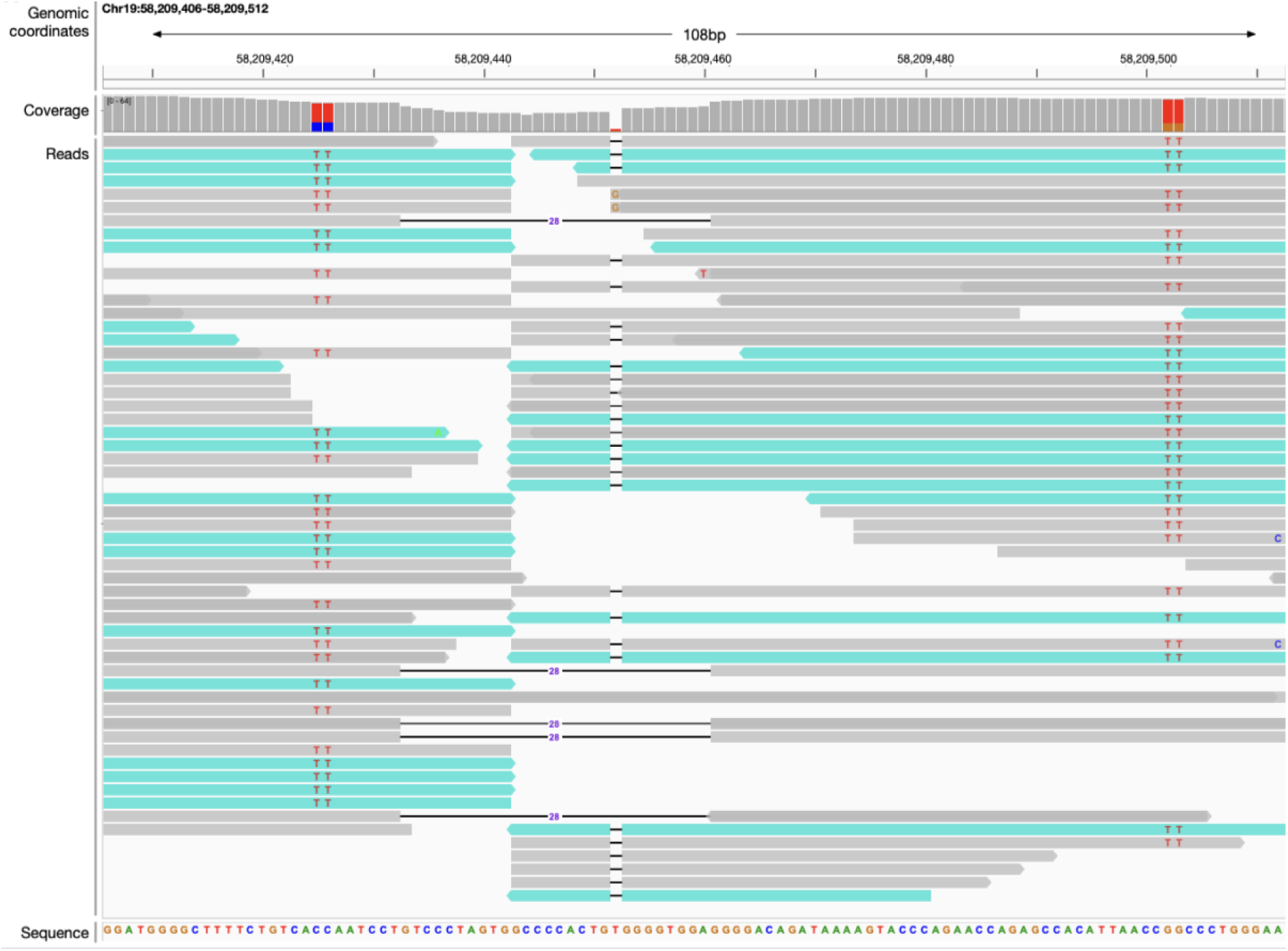
Detection of TetOn3G-FRT-BSD^R^ insertion into *AAVS1* locus. Related to Figure 2. Integrative Genomic Viewer (IGV) plot automatically generated by SeqVerify analysis of RPE1 *AAVS1* FRT8 whole genome sequencing data showing TetOn3G-FRT-BSD^R^ insertion in the *AAVS1* locus in chromosome 19. Non-chimeric reads (grey) have both ends mapped to the reference genome whereas chimeric reads (blue) have one end mapping the reference genome and another end mapping the cassette. The insertion point where chimeric reads stop aligning to the reference genome is delineated in black. TT point mutations intentionally introduced into the *AAVS1* homology arms of the insertion cassette are highlighted in red. Non-chimeric reads continuing uninterrupted through the insertion site indicate that *AAVS1* locus is intact in these reads, confirming that the insertion is heterozygous.

**Figure S3.**
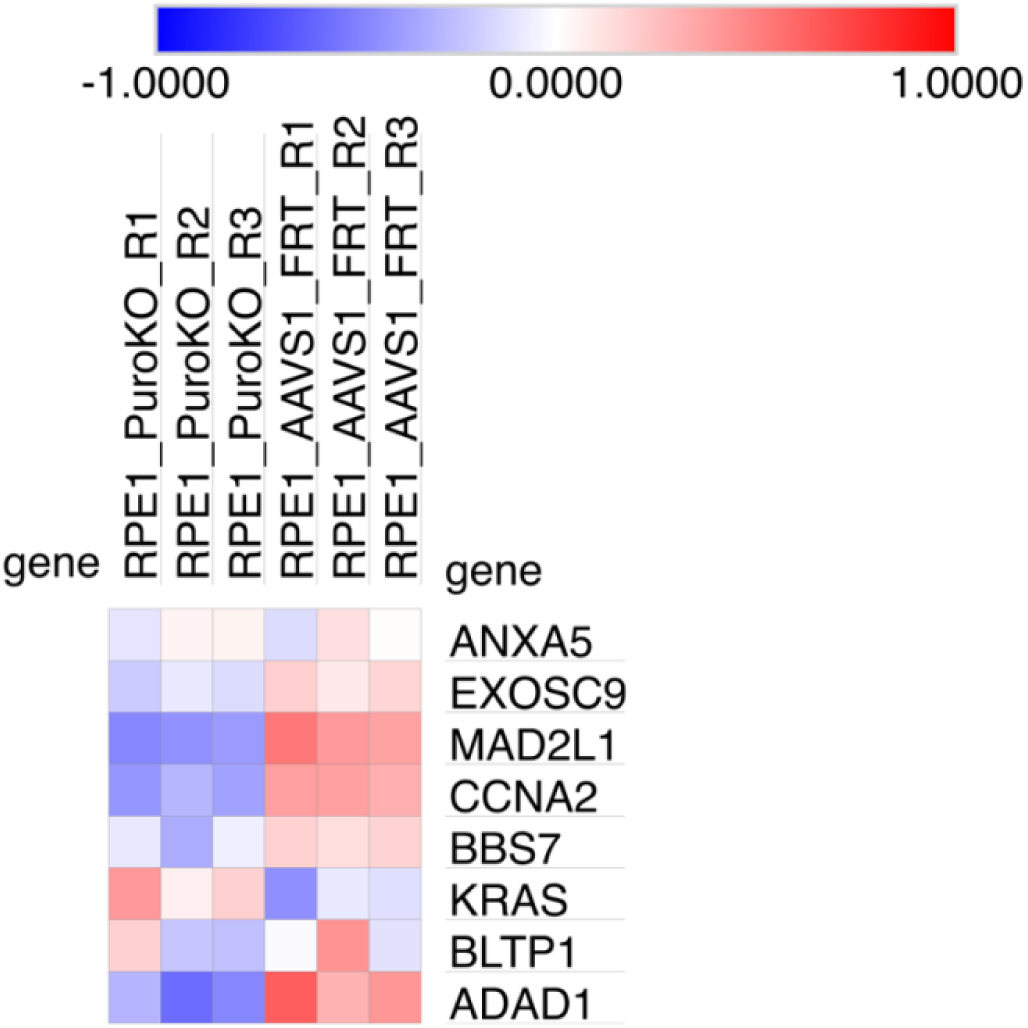
Heatmap of log2 normalised protein abundances from quantitative mass spectrometry. Related to Figure 3. The RPE1 puromycin knockout (RPE1_PuroKO_R1/2/3) and RPE1 *AAVS1* FRT8 (RPE1_AAVS1_FRT_R1/2/3) cell lines are plotted with three replicates each. The displayed genes are duplicated in whole genome sequencing and detected in quantitative mass spectrometry. Protein abundances were row-normalised across the six samples shown, such that the mean log₂ abundance for each protein was zero. Red indicates abundance above the mean and blue indicates abundance below the mean across these six samples.

**Figure S4.**
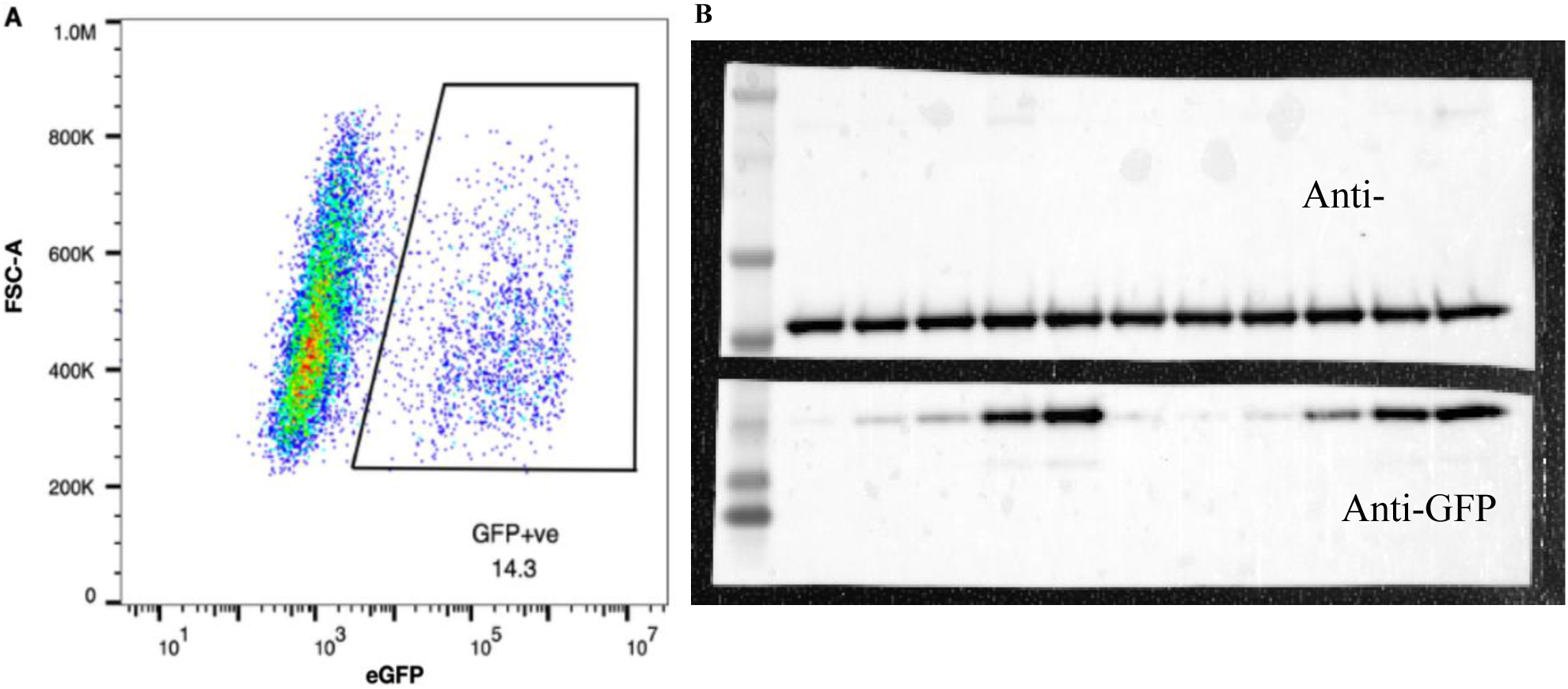
Functional validation of the FRT site and Tet-On 3G inducible system at the *AAVS1* locus. Related to Figure 4. A) Fluorescence-activated cell sorting analysis of eGFP-positive cells in RPE1 *AAVS1* FRT8 cell line expressing TRE3GS-eGFP-FRT-Neo^R^ cassette after a 16-h induction with 1 μg/ml doxycycline. B) Original, uncropped membrane corresponding to the Western blot shown in Fig. 4F. The full membrane is shown, including all lanes and molecular mass markers.

## Supplementary tables

**Table S1.**
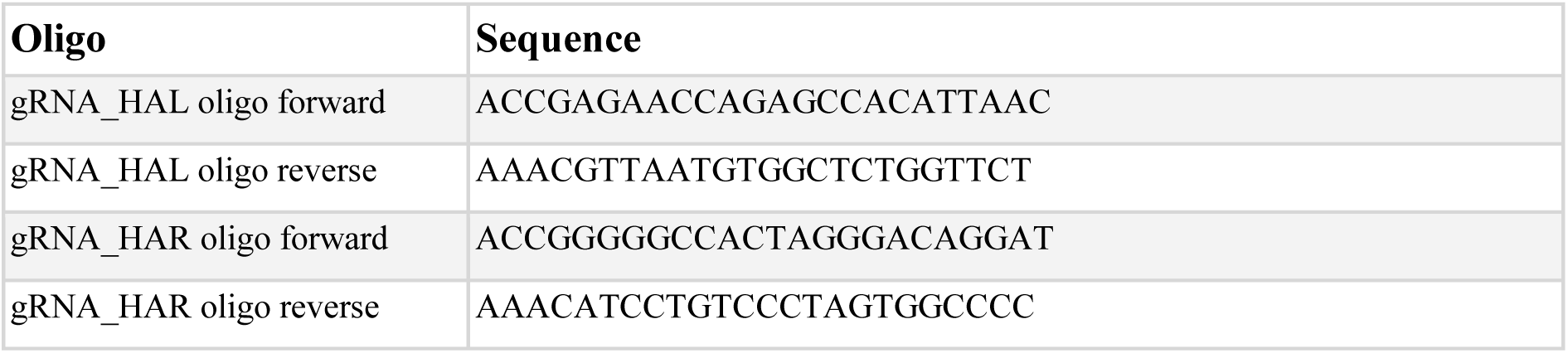
Guide RNAs targeting the AAVS1 safe harbour locus in RPE1 cells.

**Table S2.**
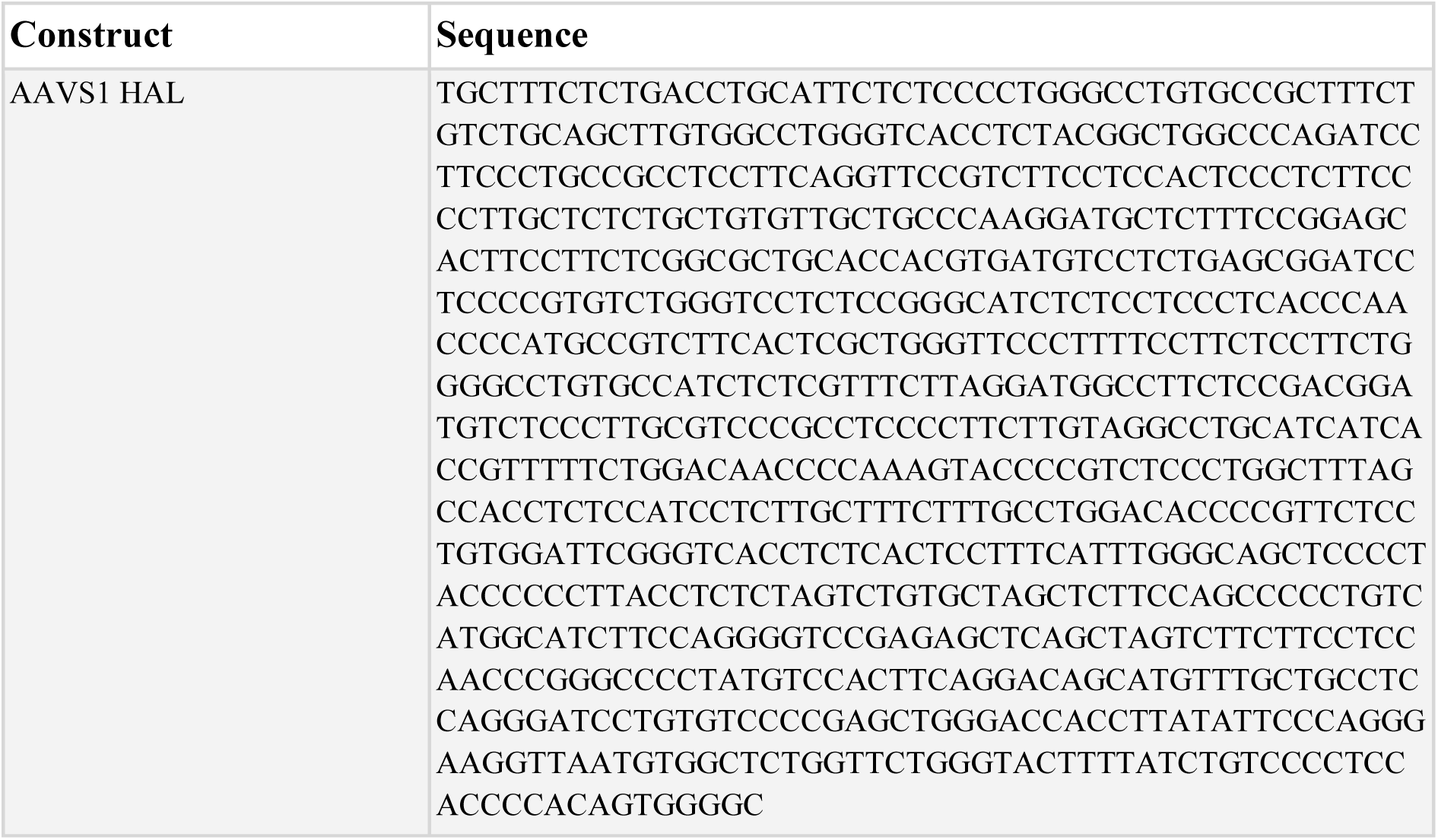
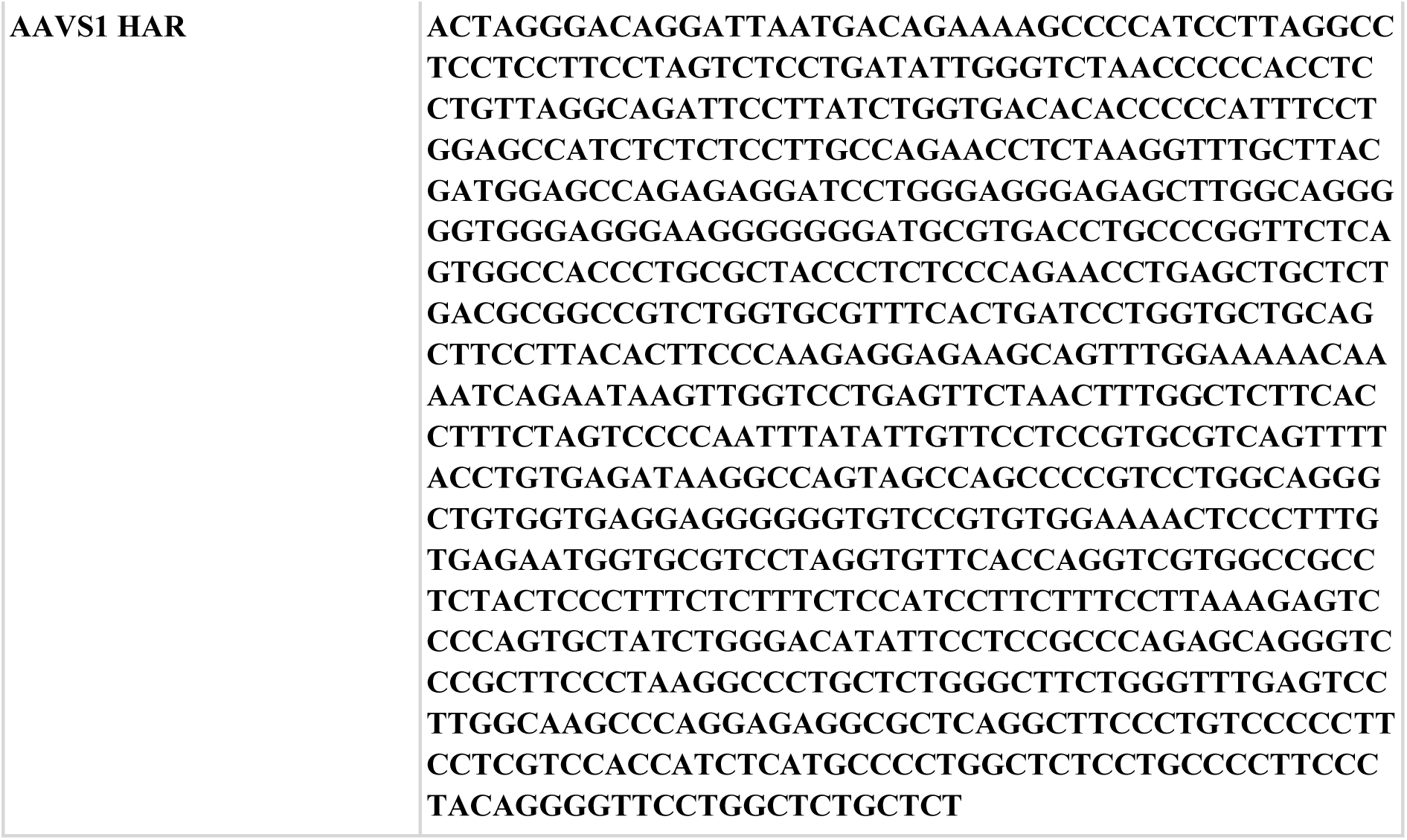
AAVS1 homology arms included in the donor plasmid for specific integration into the AAVS1 locus.

**Table S3.**
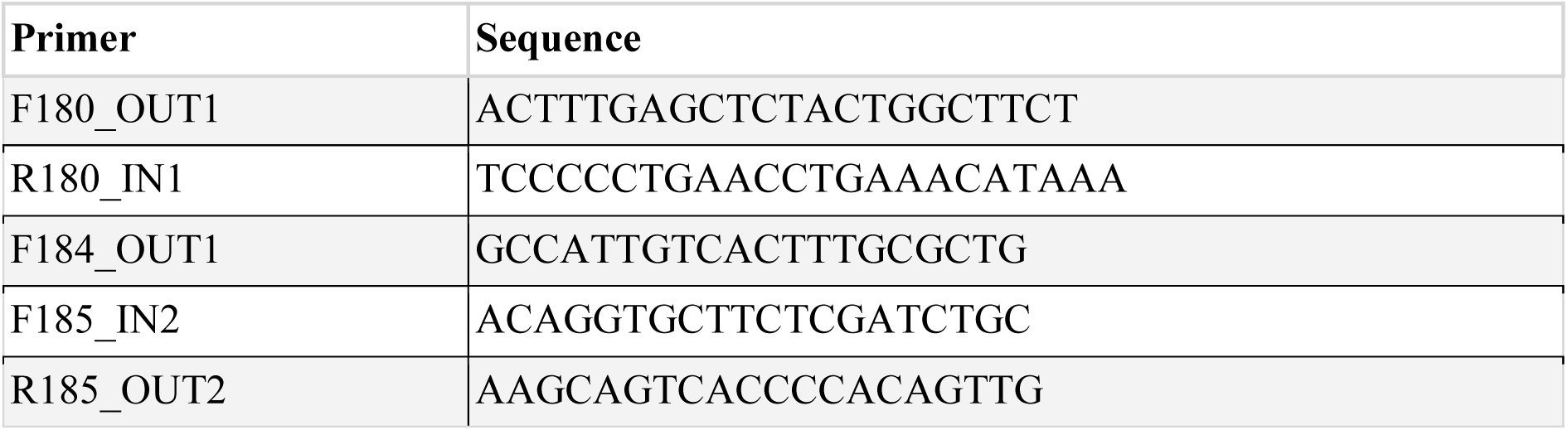
Primers used for AAVS1 locus PCR genotyping in RPE1 cells.

